# The olfactory bulb contributes to the adaptation of odor responses: the input-output transformation

**DOI:** 10.1101/829531

**Authors:** Douglas A. Storace, Lawrence B. Cohen

## Abstract

While humans and other animals exhibit adaptation to odorants, the neural mechanisms involved in this process are incompletely understood. One possibility is that it primarily occurs as a result of the interactions between odorants and odorant receptors expressed on the olfactory sensory neurons in the olfactory epithelium. In this scenario, adaptation would arise as a peripheral phenomenon transmitted into the brain. An alternative possibility is that adaptation occurs as a result of processing in the brain. Here we asked whether the olfactory bulb, the first stage of olfactory information processing in the brain, is involved in perceptual adaptation. Multicolor imaging was used to simultaneously measure the olfactory receptor nerve terminals (input) and mitral/tufted cell apical dendrites (output) that innervate the olfactory bulb glomerular layer. Repeated odor stimulation of the same concentration resulted in a decline in the output maps, while the input remained relatively stable. The results indicate that the mammalian olfactory bulb participates in olfactory adaptation.

## Introduction

To distinguish new stimuli embedded in noisy environments, humans and other animals must be able to segment novel information from the background (Linster et al., 2007; Gottfried, 2010). In olfaction, this function could be mediated by adaptation, the reduced response to repeated or continuously presented stimuli. While humans and other animals habituate to odor stimuli (Pellegrino et al., 2017), the neural mechanisms involved in this process of perceptual adaptation are incompletely understood. One possibility is that it occurs as a result of the interactions between odorants and odorant receptors expressed on the olfactory sensory neurons in the olfactory epithelium. In this scenario, adaptation would arise as a peripheral phenomenon transmitted into the brain. An alternative possibility is that adaptation occurs as a result of processing in the brain. Here we asked whether the olfactory bulb, the first stage of olfactory information processing in the brain, is involved in perceptual adaptation.

The olfactory bulb receives the projections from the peripheral olfactory sensory neuron input. All of the olfactory sensory neurons expressing the same odorant receptor converge into 1-2 glomeruli, which contain the apical dendrites of ~20-50 mitral/tufted output cells. Each mitral/tufted cell sends the transformed bulb output to a surprisingly large number of brain regions (Igarashi et al., 2012). The output is determined by a complex synaptic network that includes more than 20 interneuron types (Parrish-Aungst et al., 2007), cortical feedback (Rothermel and Wachowiak, 2014), and neurogenesis (Lledo and Saghatelyan, 2005; Lledo et al., 2006). The number of identified interneuron types of likely to increase when newer methods for determining the number are used (Parrish-Aungst et al., 2007; Shekhar et al., 2016; Zeng and Sanes, 2017). Despite its position as the first stage of olfactory information processing, the olfactory bulb has been implicated in diverse sensory functions including concentration invariance (Cleland et al., 2007; Banerjee et al., 2015; Sirotin et al., 2015; Storace and Cohen, 2017), metabolism (Fadool et al., 2011; Thiebaud et al., 2014; Riera et al., 2017), learning (Kass et al., 2013), and context-dependent processing (Doucette et al., 2011; Nunez-Parra et al., 2014; Li et al., 2017; Koldaeva et al., 2019; Wang et al., 2019).

Determining whether a brain region is involved in a neural function can be inferred by measuring its input-output transformation (Wilson et al., 2004; Storace and Cohen, 2017). Here the glomerular input and output odor activation patterns were measured and compared in response to repeated odor stimulation. In some experiments input and output were measured simultaneously in awake mice using epifluorescence imaging. In other experiments input and output were measured in separate, anesthetized preparations using 2-photon imaging. Remarkably, the olfactory sensory neuron input adapted significantly less than the mitral/tufted cell output. The results show that the bulb input maintains a relatively stable representation of the external environment, while bulb processing results in output adaptation. Thus, the olfactory bulb contributes to sensory adaptation, a function that could be important for segmenting novel odor stimuli from a static background.

## Methods

### Surgery and imaging in adult mice

All experiments were performed in accordance with relevant guidelines and regulations, including a protocol approved by the Institutional Animal Care and Use Committees at Yale University. Thy1-GCaMP6f GP5.11 transgenic mice were acquired from Jax (Stock #024339) (Dana et al., 2014). Tbx21-Cre (Jax #024507) and OMP-Cre (Jax #006668) (Li et al., 2004) transgenic mice were acquired from Jax and crossed to a floxed GCaMP6f reporter transgenic mouse (Jax #024105) (Madisen et al., 2015). OMP-GCaMP3 (Jax #029581) transgenic mice were a gift from Tom Bozza. OMP-GCaMP3 was crossed to Tbx21-Cre to create a new line in which GCaMP3 was expressed in the olfactory sensory neurons, and cre recombinase was expressed in mitral/tufted cells (OMP-GCaMP3 × Tbx21-Cre). In this line the output cells could be targeted with cre-dependent AAVs. Animals used in the study were confirmed to express the appropriate transgenes via genotyping by Transnetyx (Cordova, TN)

For all surgical procedures, male or female adult (40 – 100 days old) mice were anesthetized with a mixture of ketamine (90 mg kg^−1^) and xylazine (10 mg kg^−1^). Anesthesia was supplemented as needed to maintain areflexia, and anesthetic depth was monitored periodically via the pedal reflex. Animal body temperature was maintained at approximately 37.5 C° using a heating pad placed underneath the animal. For recovery manipulations, animals were maintained on the heating pad until awakening. Local anesthetic (1% bupivacaine, McKesson Medical) was applied to all incisions. Respiration was recorded with a piezoelectric sensor.

Calcium dye (Cal-590 Dextran, #20509, AAT Bioquest, Sunnyvale, CA) was loaded into olfactory sensory neurons of mice via the time-honored, yet thorny approach described by Wachowiak and Cohen (2001). Mice were anesthetized, placed on their back, and an 8 μl mixture of 4%/0.2% calcium dye/Triton-X was drawn into a Hamilton syringe with a flexible plastic tip, which was inserted ~10 mm into the nasal cavity. 2 μl of the dye/triton mixture was infused into the nose every 3 minutes. Mice were allowed to recover for at least 4 days prior to optical measurements. The olfactory sensory neurons were also measured using the genetically encoded calcium indicators GCaMP3 and GCaMP6f (Tian et al., 2009; Chen et al., 2013). The mitral/tufted cells were measured using GCaMP6f, jRGECO1a, jRCaMP1a and ArcLight (Jin et al., 2012; Chen et al., 2013; Dana et al., 2014; Dana et al., 2016). GCaMP6f was endogenously expressed in mitral/tufted cells in Thy1-GCaMP6f 5.11 and Tbx21-Cre × Flex-GCaMP6f transgenic mice. Cre-dependent adeno associated viral vectors were used to express jRGECO1a, jRCaMP1 and ArcLight in Tbx21-Cre transgenic mice.

For epifluorescence imaging, a custom head-post was attached to the top of the posterior skull using metabond (Parkell, Edgewood, NY). The skull above the dorsal olfactory bulb was thinned and covered with an optically transparent glue and allowed to dry. The skull was covered with Kwik-Sil (WPI Sarasota, FL), and the mouse was allowed to recover for 3 days. The recording chamber was a conical vial with the sealed end cut open in which the mouse was allowed to acclimate for a minimum of 3 days prior to imaging experiments. Mice were initially placed in the vial for periods of up to 10 minutes, which gradually increased up to 30 minutes. Animals did not exhibit signs of distress or prolonged struggle. During these acclimation periods, we measured odor-evoked responses in a small number of trials to confirm that expression was sufficient to detect functional signals.

Epifluorescence imaging was done using a custom microscope made out of Thorlabs (Newton, NJ) components with two Prizmatix LEDs (UHP-T-LED-455 and UHP-T-LED-545), which were filtered with a 488/10 nm (Semrock FF01-488/10), and a 575/5 nm (Semrock FF01-575/5) excitation filter, respectively. A beam combiner dichroic cube directed the light from both LEDs towards a beamsplitter dichroic mirror (Chroma 59009bs) that reflected the excitation light towards the preparation, while transmitting the fluorescence emission of both fluorophores towards the emission pathway. The fluorescence emission passed through a 175mm focal length lens (Edmund Optic), and a dual band-pass filter (Chroma 59009m) before being recorded with a NeuroCCD-SM256 camera with 2×2 (128×128 pixels) or 3×3 (84×84 pixels) binning at frame rates between 25-50 Hz using NeuroPlex software (RedShirtImaging, Decatur, GA).

LED strobing was performed by sending a trigger that defined the onset of each camera frame to an Arduino Mega 2560 which was programmed to send alternating triggers to the two LED power supplies (**Supplementary Code 1**). The precision of the Arduino board was measured using an oscilloscope (WaveAce 102, Teledyne LeCroy). The delay from detection of a triggered pulse on the camera pin to the output of either of the LED pins was between 4-25 μs. The jitter of the output of the LED pin was similarly small. The board was programmed to include a short buffer (500 μs) at the beginning and end of each camera frame during which the LED was off, to avoid the possibility of crosstalk across frames. The strobing control analysis in **Fig. 2c-d** was performed in a transgenic mouse that had GCaMP6f expressed in the olfactory sensory neurons (OMP-Cre × Ai95.flex.GCaMP6f). In this experiments odor-evoked signals were measured in one trial while the light was on continuously. In another trial, the blue light was strobed so that it was only on during alternate camera frames. The traces in **Fig. 2d** are from single trials. Similar results were observed in 2 additional preparations.

The procedures for *in vivo* 2-photon imaging were similar to those for epifluorescence. For 2-photon imaging experiments, the skull was removed and replaced with agarose and a cover glass window. All imaging took place while the animal was anesthetized on the same day as the surgery. 2-photon imaging was performed with a modified Sutter moveable objective microscope (MOM, Sutter Instruments) equipped with an 8 kHz resonant scanner (Cambridge Technologies). The tissue was illuminated with 940-980 nm laser light for GCaMP, or 1040nm for jRGECO1a using a Coherent Discovery laser (Coherent). Imaging was performed using either a Nikon CFI APO LWD 25× 1.10 N.A. a Nikon CFI LWD 16× 0.80 N.A., a Leica 20× 1.0NA, a Olympus 10× 0.6NA, a Zeiss 10× 0.5NA, or a Olympus 20× 1.0NA objective. Fluorescence emission passed through a 510 / 84 bandpass filter and was detected with a GaAsP PMT (Hamamatsu). Power delivered to the sample ranged from 75-140 mW as determined using a power meter (Thorlabs PM100D) placed underneath the microscope objective at the beginning of each experiment.

### Data Analysis

Odorant-evoked signals were collected in consecutive trials separated by a minimum of 3 minutes. The individual trials were manually inspected, and occasional trials with obvious motion artifact were discarded. Individual glomeruli were visually identified in the average fluorescence intensity, or via a frame subtraction analysis that identified activated glomeruli (Wachowiak and Cohen, 2001). For functional analyses, 2-photon movies were spatially binned to 128×128 pixels. Response amplitudes were calculated as the difference between the 0.5-2 s average around the peak of the response, and the 2 s preceding the stimulus. Fluorescence signals were converted to ΔF/F by dividing the spatial average of the pixels containing each identified glomerulus by the resting fluorescence. For epifluorescence the resting fluorescence was defined as the average of 5 frames at the beginning of the trial. For 2-photon imaging, it was defined as the average of the 3 seconds prior to the first odor presentation for each trial.

For the scaled subtraction analyses (e.g., **Fig. 2b-c**; **Fig. 3**), the pixels surrounding an activated glomerulus were selected as the “surround”. We made an effort to select nearby pixels that did not overlay glomeruli appearing in the activity map. The example traces shown as “scaled to surround” in **Fig. 2,** and **Supplementary Figs 2-4** were generated using the scaled subtraction function in Neuroplex. The values used in the quantification (e.g., **Fig. 2c**, **Fig. 3**) were generated by subtracting the difference between the center and surround values at the peak of the odor response. The responses to the first and third odor response were then normalized to the response amplitude evoked by the first odor presentation. The 2-photon data was not analyzed using the scaled subtraction analysis since 2-photon imaging is influenced less by out of focus fluorescence. The traces included in **Fig. 2** and **Supplementary Figs. 2-3** are single trial measurements. The traces in **Supplementary Fig. 4** are from aligned averages. The output traces in **Fig. 4** are from a single trial, the input traces are from an average of 4 single trials (the respiration trace is from one of the single trials). The average fluorescence intensity image in **Fig. 4a** was generated by taking the average of the entire imaging trial. All activity maps (e.g., **Fig. 2a**, **Fig. 4a**), and quantitative measurements (e.g., **Fig. 2c**, **Fig. 3** and **Fig. 4c-d**) were typically measured in averaged trials. The 2-photon recordings in **Fig. 4** were collected with the Olympus 10x 0.6NA objective.

The population analysis in **Fig. 3b** and **Fig. 4c** averaged all normalized measurements for each preparation. For the 2-photon concentration analysis in **Fig. 4d**, the normalized response to the second, third and fourth odor presentation were grouped across all preparations for input and output respectively (**Fig. 4d**). All statistical comparisons were performed using a Mann-Whitney U-test (ranksum in MATLAB). Similar levels of statistical significance were measured using a two-sample t-test (ttest2 in MATLAB), and a kruskal-wallis test (kruskalwallis in MATLAB). For 2-photon imaging experiments, up to five odorant pulses were delivered in the same trial at the same interstimulus interval in a subset of the preparations. The results were not qualitatively different from measurements with only four odor pulses. However, since this was not done in all the experiments, only the first four odor pulses were analyzed.

### Odorant stimuli and delivery

Odorants (Sigma-Aldrich) were diluted from saturated vapor using a flow dilution olfactometer (Vucinic et al., 2006). Odorants were delivered at different concentrations ranging from 0.1-6% of saturated vapor. Ethyl tiglate, methyl valerate, isoamyl acetate 2-heptanone and a mixture of 3 of the odorants were used. A photo-ionization detector (200B mini-PID, Aurora Scientific, Aurora, ON) confirmed the time course and relative concentrations delivered by the olfactometer at the beginning of experiments. The photo-ionization measurements in **Supplementary Fig. 6** were measured while the sensor was placed in front of the olfactometer located in front of the animal’s nose in a subset of the experiments.

### Histology

Mice were euthanized with euthasol, and their brains were dissected and left in 4% paraformaldehyde for a minimum of 3 days. Olfactory bulbs were embedded in 3% agarose, and cut on a vibratome in 50-75 μm thick coronal sections. Sections were mounted and coverslipped with VECTASHIELD Mounting Medium with DAPI (Vector Labs, H-1500) or Propidium Iodide (Vector Labs, H-1300). Slides were examined using a Zeiss LSM-780 confocal microscope (Carl Zeiss Microsystems). Appropriate targeting of the sensors was confirmed by visual inspection. The histological image in **Fig. 1a** is from a Thy1-GCaMP6f 5.11 transgenic mouse that expresses GCaMP6f in mitral/tufted cells, and in which the olfactory sensory neurons were labeled with Cal-590 dextran. The histological image in **Supplementary Fig. 1a** is from a Tbx21-Cre × Ai95.Flex.GCaMP6f transgenic mouse that expresses GCaMP6f in mitral/tufted cells. The histological image in **Supplementary Fig. 1b** is from an OMP-GCaMP3 × Tbx21-Cre transgenic mouse that expresses GCaMP3 in the olfactory sensory neurons, and received a cre-dependent AAV injection that expressed jRGECO1a in the mitral/tufted cells. The histological image in **Supplementary Fig. 1c** is from a Tbx21-Cre transgenic mouse that received a cre-dependent AAV that expressed ArcLight.

**Figure 1:**
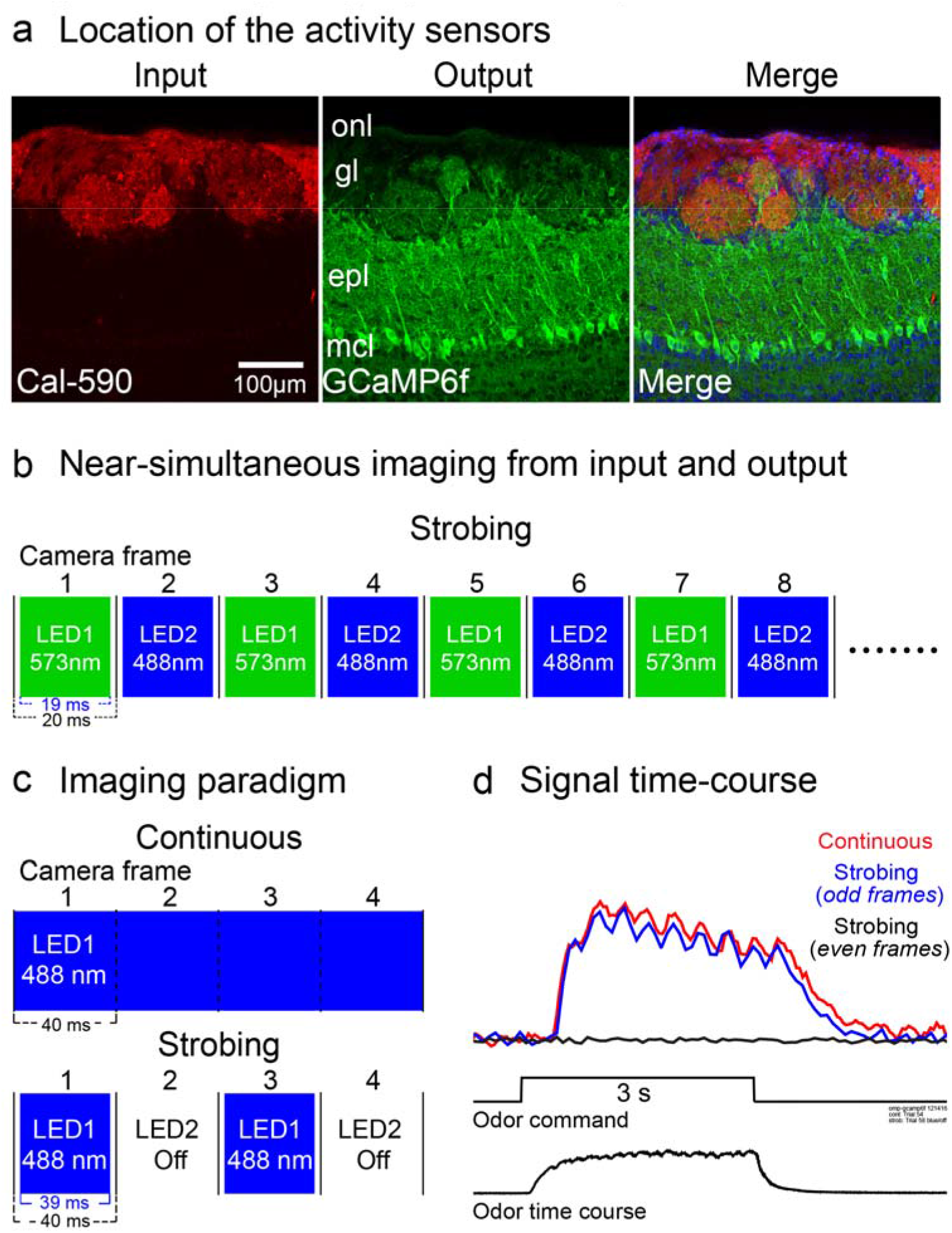
Targeting input and output. Approach for measuring the olfactory bulb input-output transformation near-simultaneously. (**a**) Targeting of spectrally distinct sensors to input and output. Histology is from a transgenic mouse expressing GCaMP6f in the mitral/tufted cells (Thy1-GCaMP6f 5.11) in which the olfactory sensory neurons were labeled with the organic calcium dye Cal-590 dextran via nasal infusion. (**b**) Paradigm for near-simultaneous imaging from the two fluorophores. (**c-d**) Multiplexing LEDs does not notably impact signal-to-noise ratio. In one experiment GCaMP6 was imaged continuously (**c**, *top*), or was multiplexed so that alternate frames were blank (**c**, *bottom*). (**d**) Odor-evoked fluorescence signals measured from the illuminated frames were not distinguishable from the continuous recording, and no light was detectable in the blank camera frames (black trace). Similar results were obtained in 2 other preparations.

**Figure 2:**
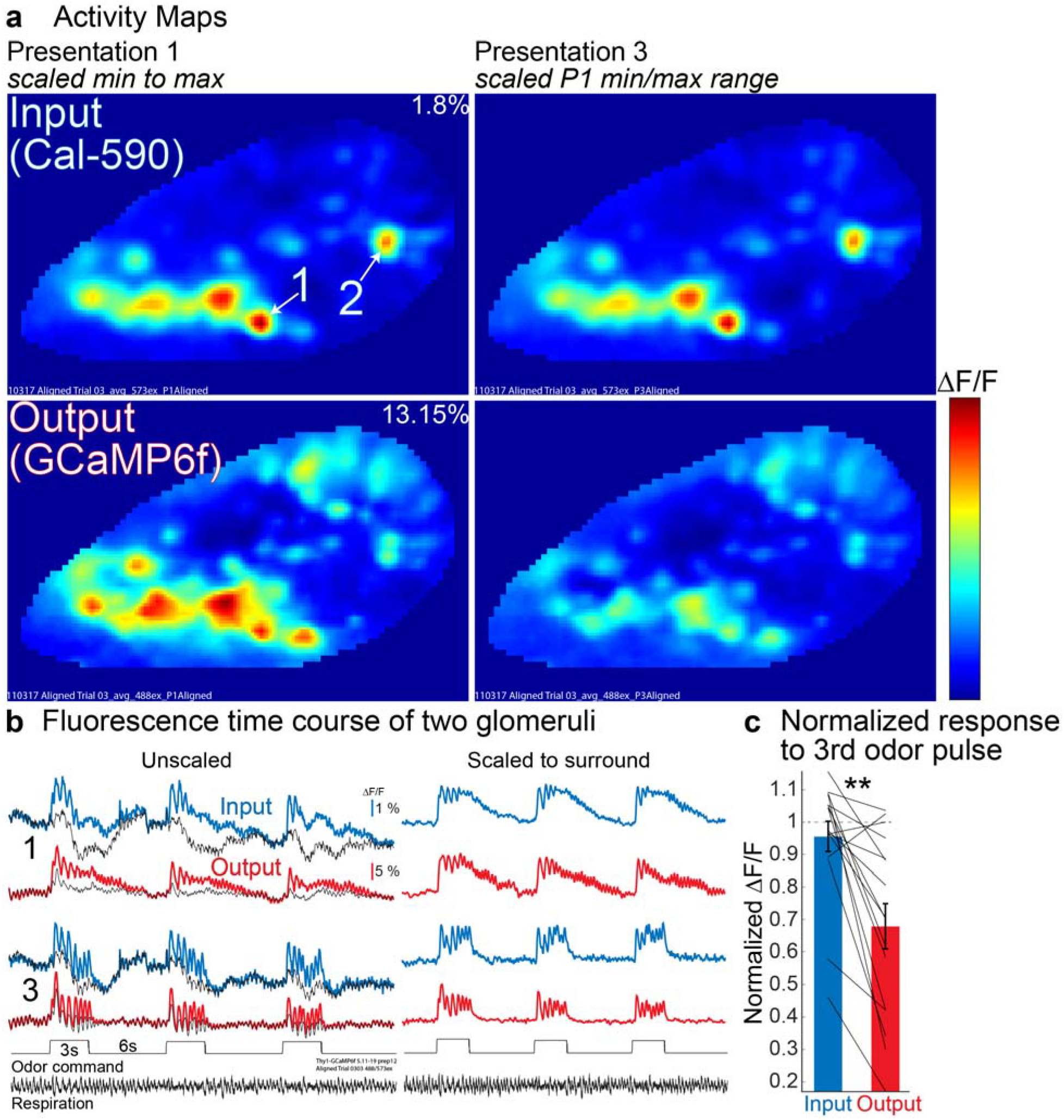
The output adapts more than the input. The bulb output adapts more than the input in response to repeated odor stimulation in a Thy1-GCaMP6f 5.11 transgenic mouse that had olfactory sensory neurons loaded with Cal-590 dextran. (**a**) Input (*top*) and Output (*bottom*) activity maps are shown in response to the first odor presentation scaled to their minimum and maximum ΔF/F values (*left panel,* max ΔF/F indicated on panel). The activity map evoked by the third odor pulse is shown with the color scale set to the ΔF/F maximum value evoked by the first odor pulse (*right panel*). (**b**) Fluorescence time course of the near-simultaneously measured input and output for 2 glomeruli from one of the single trials used to generate the activity maps in panel **a**. *Unscaled*: The colored traces are from the center activated glomeruli. The signal from the immediate surround is shown as the superimposed thin black trace. *Scaled to surround*: Fluorescence traces that have had the surround subtracted from the glomerular center to remove common noise. (**c**) Quantification of the signal size of the 3^rd^ odor presentation normalized to the amplitude evoked by the 1^st^ odor presentation for 16 glomeruli. The dashed line indicates a measurement that would reflect zero adaptation. The traces in panel **b** are from single trials low-pass filtered at 4 Hz. The maps in panel **a** and the quantification in **c** are from aligned averages. The odor used was 2% saturated vapor methyl valerate. ** p < 0.005.

**Figure 3:**
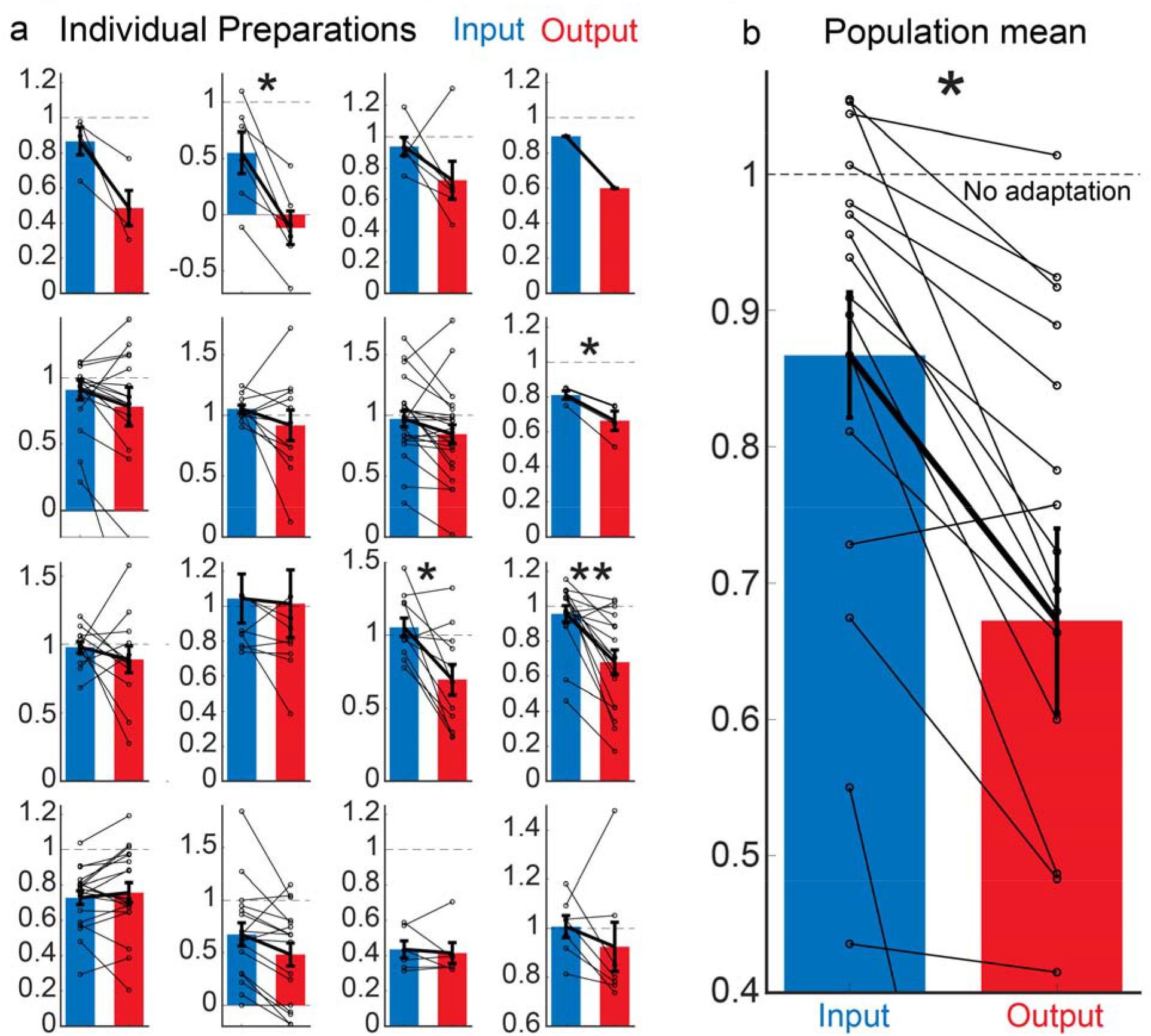
Normalized response to 3rd odor pulse. Population summary of individual preparations in which both input and output were measured in the same hemibulbs of awake mice. Measurements are shown as the response to the third odor pulse normalized to the ΔF/F amplitude evoked by the first. (**a**) Signal size measurements from identified glomeruli that were present in both input and output for 16 preparations. Lines show the direction of the same glomeruli for input and output. (**b**) Quantification of signal size across the population of all 16 preparations. Between 1-23 glomeruli were measured in each preparation (10.6 ± 1.6). Input was measured using Cal-590 dextran or GCaMP3 in 11 and 5 preparations, respectively. Output was measured using GCaMP6f, jRGECO1a, or jRCaMP1a in 11, 3 and 2 preparations, respectively. Statistical comparisons were measured using a Mann-Whitney *U* test, * < 0.05, ** < 0.01.

**Figure 4:**
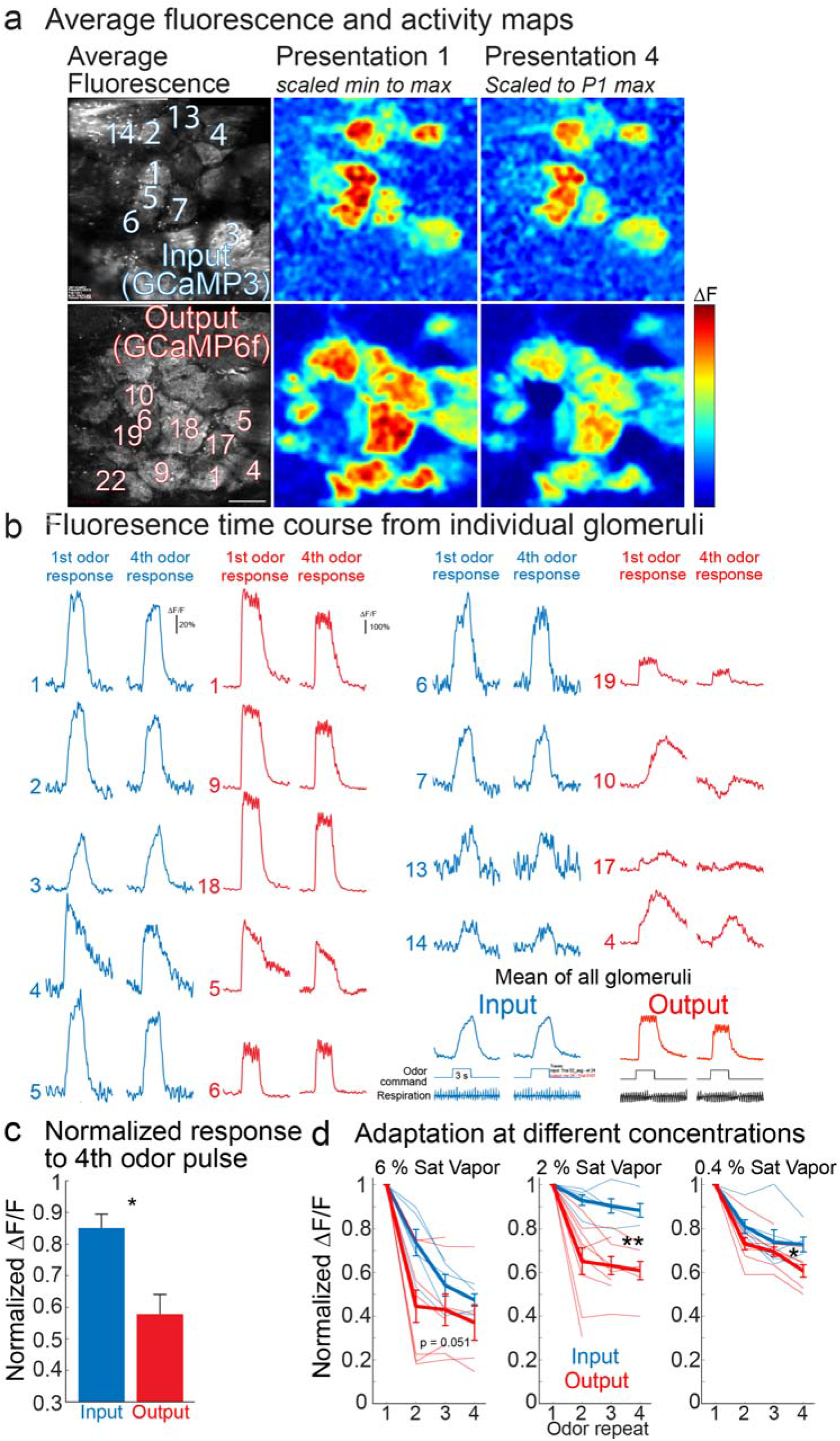
2-Photon imaging from input and output in different, anesthetized preparations. Adaptation of input and output responses measured in different, anesthetized preparations using 2-photon imaging. (**a**) Average fluorescence intensity and activity maps from two different preparations in which input (GCaMP3) or output (GCaMP6f) were measured separately. (**b**) Fluorescence time course from activated glomeruli measured in the example preparations in **a**. The response to the first and fourth odor presentation are cropped and displayed next to one another. (**c**) Quantification of the signal size of the fourth odor presentation normalized to the amplitude evoked by the first odor presentation for all of the glomeruli identified in these two recording trials. (**d**) Population summary of adaptation of input and output at three different concentrations. Each line is from a different preparation. The input and output preparations were measured in response to ethyl tiglate (2% saturated vapor) and methyl valerate (2% saturated vapor), respectively. Scalebar in panel **a**, 100 μm.

The fluorescence for the images in **Fig. 1a** and **Supplementary Fig. 1** were detectable without additional amplification steps, were contrast-enhanced and sharpened (unsharp mask, both applied uniformly to entire image), and were cropped and pseudo-colored using Zen Lite 2011 (Carl Zeiss Microsystems), Adobe Photoshop and Illustrator (Adobe Systems Inc.).

## Results

### The method for simultaneous measurements of input and output

A more efficient version of our approach to measure the olfactory bulb input-output transformation was developed (Storace and Cohen, 2017). This strategy involves using spectrally separated optical indicators to near-simultaneously measure the signals from the olfactory sensory neuron input and mitral/tufted cell output. Olfactory sensory neurons were loaded with an organic calcium dye via nasal infusion (Friedrich and Korsching, 1997; Wachowiak and Cohen, 2001; Fried et al., 2002) in a transgenic mouse that expressed a genetically encoded calcium indicator (GECI) in the mitral/tufted output cells (**Fig. 1a**). Input and output measurements were made using a custom microscope with two LED light sources that were filtered to the peak excitation wavelength of the input or output fluorophore. The light from both LEDs passed through a beamsplitter dichroic mirror which allowed the excitation light to reach the preparation, and the fluorescence emission to reach the camera. The LEDs were synchronized with the camera so that only one LED illuminated the preparation per camera frame (**Fig. 1b**) (Miyazaki and Ross, 2015, 2017; Miyazaki et al., 2018). Because the excitation spectra of the fluorophores used in these experiments had minimal overlap, this approach reduced cross-talk between the fluorophores.

Strobing LEDs did not noticeably impact the imaging signal-to-noise ratio. This can be seen by comparing measurements from trials using continuous illumination to those in which the LED is strobed, in a preparation that only had a single fluorophore (GCaMP6f). Odor-evoked signals were measured during trials in which the blue LED was on continuously, and in separate trials where the blue LED was only on during alternate camera frames (**Fig. 1c,** *top vs bottom*). The signals from the continuous imaging trial were similar to those from the LED-On frames of the strobing trial, while the LED-Off frames were blank (**Fig. 1d,** *black trace*). Thus, this approach is suitable for near-simultaneous measurements of the olfactory bulb input-output transformation.

### Simultaneous measurements from the same input and output glomeruli in the same mouse (awake)

Odor-evoked responses were measured from the bulb input and output in response to repeated presentations of the same odor stimulus (three three second presentations with a six second interstimulus interval) in awake head-fixed mice. In one variation of the experiment, measurements were made in the Thy1-GCaMPf 5.11 transgenic line that expresses GCaMP6f in the mitral/tufted cells (Dana et al., 2014; Iwata et al., 2017) (**Fig. 2)**. The response to each odor presentation was visualized by performing a frame subtraction analysis for both the input and output (**Fig. 2a**). Single trial fluorescence time course measurements from two activated glomeruli are shown in **Fig. 2b**.

In this preparation the input measurements were notably stable across odor repeats, and exhibited minimal adaptation. The activity map evoked by the third odor presentation was very similar to the activity map evoked by the first odor presentation. This is visualized by scaling the max ΔF/F value of the first activity map to the max ΔF/F of the first activity map (**Fig. 2a**, *input, left versus right panel*). In contrast, the magnitude of the output response changed substantially across odor presentations. Applying the same scaling analysis to the output activity maps revealed reductions in signal amplitude in most of the active glomeruli (**Fig. 2a**, *output, right panel*).

The presence of a peak in the activity map indicates that a glomerulus is more strongly activated than the surround. This is evident by comparing the fluorescence time course measured from the center of a glomerulus to its surrounding area (**Fig. 2b**, *color versus black traces*). For the input measurements, the separation between the center and surround were similar for each odor pulse. In contrast, the separation between the center and surround of the output glomeruli became smaller (**Fig. 2b**, *output*). The results from these two glomeruli are consistent with the changes visualized in the input and output activity maps, respectively (**Fig. 2a** vs **2b**). The difference between the center and surround of activated input and output glomeruli was used to quantify the amount of adaptation that occurred as a function of odor repeats. This analysis removed global noise and demonstrates that the amplitude of the bulb input declined much less than the output (**Fig. 2b**, *scaled to surround*). The amplitudes of the scaled glomerular responses were normalized to the response evoked by the first odor presentation. For this preparation, the input declined to 0.96 ± 0.05, while the output declined to 0.68 ± 0.07 (**Fig. 2c**) (p < 0.005; N = 16 glomeruli).

The same experiment was performed in the Tbx21-Cre transgenic mouse line that expresses cre recombinase in mitral/tufted cells. In one variation of the experiment, GCaMP6f was targeted to the mitral/tufted by means of a cross to a transgenic GCaMP6f reporter mouse (Tbx21-Cre × Ai95-Flex-GCaMP6f). Histology from one of these mice showing GCaMP6f targeted to the mitral/tufted cells is shown in **Supplementary Fig. 1a**. An example preparation is shown in **Supplementary Fig. 2**, which has an identical display arrangement to **Fig. 2**. On average the output declined more than the input (Input: 0.87 ± 0.08; Output: 0.49 ± 0.1; p =0.06; N = 4 glomeruli) (**Supplementary Fig. 2c**).

In another variation of the experiment, the Tbx21-Cre transgenic mouse line was crossed to a transgenic mouse in which GCaMP3 (Cichy et al., 2019) was expressed in the bulb input (OMP-GCaMP3 × Tbx21-Cre). The mitral/tufted cell output was targeted with either jRCaMP1a or jRGECO1a (Dana et al., 2016) using Cre-dependent adeno-associated viruses. Histology demonstrating targeting of the sensors in this double transgenic mouse line is shown in **Supplementary Fig. 1b**. jRCaMP1a was used for the preparation in **Supplementary Fig. 3** where the input and output declined to 0.97 ± 0.06 and 0.85 ± 0.07 (N = 23 glomeruli), respectively.

jRGECO1a was used for the preparation in **Supplementary Fig. 4**. However, input and output measurements performed in alternate imaging trials because blue light activates slow and sustained rises in jRGECO1a fluorescence, which confounded simultaneous measurements (**Supplementary Fig. 5**). Regardless, the results were consistent with the other preparations where the normalized response to the 3^rd^ odor presentation for input and output was 1.0 ± 0.04 and 0.92 ± 0.1 (N = 7 glomeruli), respectively.

### Summary of same glomeruli and same animal comparisons

Input and output measurements were performed in 16 preparations using 2 different input sensors (Cal-590: 11 preparations; GCaMP3: 5 preparations) and 3 different output sensors (GCaMP6f: 11 preparations, jRCaMP1a: 2 preparations; jRGECO1a: 3 preparations) (**Fig. 3a**). Not all glomeruli exhibited the same behavior in each preparation (e.g., the output declined less than the input in some glomeruli), and not all preparations had within comparisons that were statistically significant. However, most of the experiments had means in the same direction (but not all, see #13) (**Fig. 3a**). The average normalized response to the 3^rd^ odor preparation was averaged across all activated glomeruli per preparation (between 1-23 glomeruli per preparation). Overall, the input and output declined to 0.85 +/− 0.04 and 0.65 +/− 0.07, respectively (**Fig. 3b**; p < 0.02).

### Measuring adaptation using a voltage sensor

Output measurements were also performed using the genetically encoded voltage indicator (GEVI) ArcLight targeted to the mitral/tufted cells in anesthetized Tbx21-Cre transgenic mice (using a cre-dependent AAV) (Histology shown in **Supplementary Fig. 1c**). In three preparations both input and output were measured, but in opposite hemibulbs. Output alone measurements were made in five additional preparations. The differences were consistent with the measurements using calcium imaging alone, although the smaller sample size yielded a non-statistically significant difference (*not shown*).

### Olfactometer variability cannot explain the result

Because many of these measurements were performed simultaneously, the differences are unlikely to be due to variability in the animal’s state or respiration. Furthermore, we confirmed that the olfactometer delivered repeatable odor pulses using a photoionization detector (**Supplementary Fig. 6**). The amplitude of the 3^rd^ odor pulse changed less than 2% relative to the 1^st^ odor pulse (1.3 ± 1.7 % SEM) when averaging many individual trials using different odorants and concentrations. Thus, measurements from input and output using different combinations of sensors in awake and anesthetized mice show that the olfactory bulb likely to contributes to odor adaptation.

### Measuring adaptation in input and output using 2-photon imaging in separate preparations

The optical signals from wide-field imaging can be influenced by fluorescence originating from above and below the focal plane of the objective. In contrast, 2-photon microscopy has a much smaller depth-of-field (Denk et al., 1990). We used 2-photon microscopy to measure input and output to confirm that the results measured using 1-photon imaging (i.e., **Figs. 2-3 and Supplementary Figs. 2-4**) occurred in the glomerular layer. However, in these experiments input and output were measured in different, anesthetized preparations. The input signals were measured in transgenic mice expressing either GCaMP3 (OMP-GCaMP3, N = 5 preparations) or GCaMP6f (OMP-Cre × Flex-GCaMP6f, N = 1 preparation). Output signals were measured in transgenic mice expressing either GCaMP6f (Thy1-GCaMP6f 5.11, N = 5 preparation; Tbx21-Cre × Flex-GCaMP6f, N = 3 preparations) or jRGECO1a (Tbx21-Cre injected with a Cre-dependent jRGECO1a expressing AAV, N = 2 preparations).

Odor responses were measured in 103 individual input glomeruli (between 7 and 31 glomeruli per preparation). However, the same glomeruli were tested across multiple stimulus conditions (e.g., different odorant-concentration pairs) if the preparation allowed it, resulting in a total of 278 total input measurements (6% N = 100; 2% N = 165; 0.4% N = 118; methyl valerate, isoamyl acetate and 2-heptanone). Odor responses were measured in 383 individual output glomeruli (between 9 and 56 glomeruli per preparation). Recording different odor-concentration pairs yielded a total of 705 total measurements (6% N = 205; 2% N = 284; 0.4% N = 216; methyl valerate, isoamyl acetate, ethyl tiglate and 2-heptanone).

### Effects of adaptation on input versus output measured using 2-photon imaging

The activity maps and fluorescence traces from 2 preparations in which only the input or output were measured are shown in **Fig. 4**. The activity maps are scaled in the same manner as those in **Fig. 2**. Consistent with the epifluorescence imaging experiments, the input measurements were much more similar across odor repeats, while the output declined (**Fig. 4a**, see reduced hot colors for most output glomeruli). Fluorescence traces from some of the activated glomeruli are shown to demonstrate the differences (**Fig. 4b**), along with the average of all activated glomeruli (**Fig. 4b**, *mean of all glomeruli*). The mean normalized response to the fourth odor presentation for the input preparation was 0.85 ± 0.06 (N = 15 glomeruli, 2% saturated vapor, ethyl tiglate), while the mean of the output preparation was 0.57 ± 0.04 (N = 27 glomeruli, 2% saturated vapor, methyl valerate) (p < 0.05).

Next, we examined whether odorant concentration influenced adaptation of either input or output. Input and output adaptation were measured by pooling the normalized responses to the second, third and fourth odor presentation for each preparation in response to three different odor concentrations (6%, 2% and 0.4% of saturated vapor for methyl valerate, 2-heptanone, ethyl tiglate, and a mixture of the three). Both input and output were affected by odorant concentration, however the output adapted more than the input at all concentrations (6% Input: 0.56±0.03 Output: 0.34±0.08, p = 0.051; 2% Input: 0.9±0.02, Output: 0.54±0.05, p < 0.001; 0.4% Input: 0.76±0.03, Output: 0.67±0.02, p < 0.05; Mean ± SEM; Mann-Whitney *U*-Tests).

## Discussion

A long-standing question in the field is whether the olfactory bulb contributes to olfactory adaptation. Here we provide the first direct comparison between the olfactory sensory neuron input and the mitral/tufted cell output from the same preparation. Measurements from the olfactory sensory neurons were relatively stable, while the mitral/tufted signal declined in amplitude. The results demonstrate that the olfactory sensory neurons provide a faithful representation of the sensory environment, while processing in the brain reduces the output signal after repeated presentations. This function would be important for segmenting novel and potentially important odor stimuli from ongoing sensory information in the background.

### Methods consideration

#### Near-simultaneous measurements of input and output using strobing LEDs

Because adaptation is experience-dependent, we wanted a way to perform the input and output measurements simultaneously. Multiplexing the LED was preferable to using a dual band-pass filter to simultaneously excite both fluorophores since many fluorophores have broad excitation and emission spectra, which we found sometimes introduced cross-talk between the channels during preliminary testing (not shown). We concluded that the noise introduced by this approach was negligible given the 20 ms duration camera frames. Each frame was illuminated for 19000/20000 μs, so the delays and jitter we observed (< 20 μs) were ~0.1% of the baseline signal, a noise level below our baseline noise.

#### Differences between Thy1-GCaMP6f and Tbx21-Cre transgenic mice

While the Thy1-GCaMP6f line has selective mitral/tufted expression in the olfactory bulb, it also has cortical expression (Dana et al., 2014; Iwata et al., 2017). We reasoned that some of these cells could feedback to the bulb and introduce confounding fluorescence during widefield measurements. Thus, we repeated this experiment using the Tbx21-Cre mouse line, which has expression of cre recombinase exclusively restricted to the mitral/tufted cells (Mitsui et al., 2011). We measured similar results in these mice. *Sensor differences*. Different optical indicators can vary in their signal-to-noise, calcium affinity (Kd) and Hill coefficient. We think it is unlikely that our reported differences are artifactual because earlier widefield measurements were carried out with a number of different calcium and voltage indicators and they yielded similar results (Storace and Cohen, 2017; Storace et al., 2019). The sensors used in the study provide relatively a relatively linear readout of spike rate over a large dynamic range (Tian et al., 2009; Wachowiak et al., 2013; Tischbirek et al., 2015). Furthermore, the combination of input and output sensors was varied to control for differences in sensor calcium affinity and Hill coefficient (e.g., **Fig. 3**).

### Comparison with previous studies and possible mechanisms

Odorant receptors are known to exhibit adaptation to continuous odor streams or paired pulse stimulation. However, intact olfactory sensory neurons stimulated with paired pulses of odorants exhibited substantial recovery of the initial response with a 6 second delay between pulses (Zufall and Leinders-Zufall, 2000). We used an intertrial interval of six seconds which allowed time for the input signal to partly recover from any adaptation that occurred from an odor pulse.

It seems likely that the adaptation we measured involves local mechanisms within the olfactory bulb as well as processing across multiple brain areas. For example, the mitral/tufted cells send axons to more than 12 brain areas, and many of those brain areas send projections back to the olfactory bulb (Davis and Macrides, 1981; Igarashi et al., 2012). This idea is consistent with a report of adaptation measurements in neurons located in the Drosophila mushroom body (Hattori et al., 2017), a brain area analogous to the piriform cortex in mice which provides significant feedback to the olfactory bulb. Furthermore, mitral/tufted cells are experience and context-dependent (Doucette et al., 2011), indicating that the mitral/tufted cells are shaped by sensory experience.

#### Differences in presentation of output signals here and in past studies

The present study demonstrates that mitral/tufted cells sometimes exhibit suppression in response to odor presentation. This is in contrast with some prior reports of glomerular measurements from mitral/tufted cells in anesthetized mice (Fletcher et al., 2009; Ogg et al., 2015; Storace and Cohen, 2017). Anesthesia has a known dramatic effect on the physiology of mitral/tufted cells (Rinberg et al., 2006; Wachowiak et al., 2013). Thus, anesthetic state may help explain some of the differences in the output responses described here and in previous reports (Fletcher et al., 2009; Ogg et al., 2015; Storace and Cohen, 2017).

### Future work and significance

The results indicate that processing within the bulb can significantly transform the stable olfactory sensory neuron input. This function would be useful for higher brain regions to determine whether an olfactory input is novel or in the background. Future studies need to determine whether the adaptation seen in the mitral/tufted cells can be modulated. How long does it take for the mitral/tufted cells to recover? Is the adaptation specific to the odorants used in this study? Is the degree of adaptation altered by learned associations, which are known to influence mitral/tufted cell activity (Wilson and Sullivan, 1992; Doucette et al., 2011; Nunez-Parra et al., 2014; Ross and Fletcher, 2018)? What are the exact mechanisms responsible for this adaptation?

In addition, future studies are needed to determine if other olfactory perceptions are also computed in the olfactory bulb. In a previous study we showed that the olfactory bulb makes a substantial contribution to the perception of odorant concentration invariance (the ability to recognize the odorant as the same across a range of odorant concentrations) (Storace and Cohen, 2017; Storace et al., 2019). Thus, the olfactory bulb appears to be participating in two perceptual computations simultaneously. Presumably the large number of olfactory bulb interneuron types contribute to this ability (Parrish-Aungst et al., 2007).

**Supplementary Fig. 1**: Histology from two different transgenic mice showing expression of the sensors in the expected locations. (**a**) Tbx21-Cre × Ai95.Flex.GCaMP6f transgenic mouse resulted in mitral/tufted cell specific expression. (**b**) OMP-GCaMP3 × Tbx21-Cre transgenic mouse. GCaMP3 is targeted to the bulb input. jRGECO1a was expressed in the mitral/tufted cells using a cre-dependent AAV. (**c**) ArcLight is targeted to the mitral/tufted cells using a cre-dependent AAV in a Tbx21-Cre transgenic mouse.

**Supplementary Figure 2**: The bulb output adapts more than the input in response to repeated odor stimulation in a Tbx21-Cre × Ai95.Flex.GCaMP6f transgenic mouse that had Cal-590 loaded into its OSNs. (**a**) Input (*top*) and Output (*bottom*) activity maps are shown in response to the first odor presentation scaled to their minimum and maximum ΔF/F values (*left panel*, max ΔF/F indicated on panel). The activity map evoked by the third odor pulse is also shown with the color scale fixed to the ΔF/F range evoked by the first odor pulse (*right panel*). (**b**) Fluorescence time course of input and output for 2 glomeruli from one of the single trials used to generate the activity maps in panel **a**. *Unscaled*: The colored traces are from the center activated glomeruli, while the signal from the immediate surround is shown as the superimposed thin black trace. *Scaled to surround*: Fluorescence traces that have had the surround subtracted from the glomerular center. (**c**) The differences between the center and surround of each glomerulus was measured, and the signal size evoked by the third odor presentation was normalized to the amplitude evoked by the first odor presentation for 4 glomeruli. The traces in panel **b** are from single trials low-pass filtered at 4 Hz. The maps in panel **a** and the quantification in **c** are from aligned averages. The odor used was 2% saturated vapor isoamyl acetate.

**Supplementary Fig. 3**: The bulb output adapts more than the input in response to repeated odor stimulation in an OMP-GCaMP3 × Tbx21-Cre transgenic mouse that was injected with a cre-dependent AAV expressing jRCaMP1a. The data display arrangement and legend are otherwise identical to **Fig. 2**. (**a**) Input (*top*) and Output (*bottom*) activity maps are shown in response to the 1^st^ odor presentation scaled to their minimum and maximum ΔF/F values (*left panel*, max ΔF/F indicated on panel). The activity map evoked by the 3^rd^ odor pulse is also shown with the color scale fixed to the ΔF/F range evoked by the first odor pulse (*right panel*). (**b**) Fluorescence time course of input and output for 2 glomeruli from one of the single trials used to generate the activity maps in panel **a**. *Unscaled*: The colored traces are from the center activated glomeruli, while the signal from the immediate surround is shown as the superimposed thin black trace. *Scaled to surround*: Fluorescence traces that have had the surround subtracted from the glomerular center. (**c**) The differences between the center and surround of each glomerulus was measured, and the signal size evoked by the third odor presentation was normalized to the amplitude evoked by the first odor presentation for 23 glomeruli. The traces in panel **b** are from single trials low-pass filtered at 4 Hz. The maps in panel **a** and the quantification in **c** are from aligned averages. The odor used was 2% saturated vapor methyl valerate.

**Supplementary Fig. 4**: The bulb output adapts more than the input in response to repeated odor stimulation in an OMP-GCaMP3 × Tbx21-Cre transgenic mouse that was injected with a cre-dependent AAV expressing jRGECO1a. The data display arrangement and legend are otherwise identical to **Fig. 2**. (**a**) Input (*top*) and Output (*bottom*) activity maps are shown in response to the 1^st^ odor presentation scaled to their minimum and maximum ΔF/F values (*left panel*, max ΔF/F indicated on panel). The activity map evoked by the 3^rd^ odor pulse is also shown with the color scale fixed to the ΔF/F range evoked by the first odor pulse (*right panel*). (**b**) Fluorescence time course of the near-simultaneously measured input and output for 2 glomeruli from one of the single trials used to generate the activity maps in panel **a**. *Unscaled*: The colored traces are from the center activated glomeruli, while the signal from the immediate surround is shown as the superimposed thin black trace. *Scaled to surround*: Fluorescence traces that have had the surround subtracted from the glomerular center. (**c**) The differences between the center and surround of each glomerulus was measured for the first and third odor pulse and normalized to the amplitude evoked by the first odor presentation for 7 glomeruli measured in this preparation. The activity maps and traces in panels **a**-**b**are from an average of multiple single trials aligned to the beginning of the odor response, and were low-pass filtered at 4 Hz. The odor used was 2% saturated vapor methyl valerate.

**Supplementary Fig. 5**: Blue light evokes a large increase in jRGECO1a fluorescence emission in the absence of odor presentation. (**a**) The green LED (573 nm) illuminated the preparation every other camera frame. No response is present while alternate frames are blank. The introduction of blue (470 nm) light on the alternate camera frames causes a large increase in jRGECO1a fluorescence emission that slowly returned to baseline after the blue LED was removed.

**Supplementary Fig. 6**: The olfactometer delivers repeatable odor pulses. (**a**) A photoionization detector was used to measure the output of the olfactometer across three odor repeats at 10 and 2% of saturated vapor. Similar recordings were taken across 8 experiments on different days. Photoionization detector amplitude measurements taken at 10%, 2%, 0.4% and 0.13% of saturated vapor (either isoamyl acetate or methyl valerate) varied by 5.8 ± 1.8% (N = 10 measurements in 5 preparations), 0.8 ± 3.7% (N = 8 measurements in 6 preparations), 4.6 ± 2.3% (N = 6, measurements in 4 preparations) and 0.6 ± 9.4% (N = 3 measurements in 3 preparations), respectively. Differences across concentrations were not statistically different.

**Supplementary Code 1**: Arduino program used to synchronize camera with alternating LEDs. The digitalwritefast library is required to significantly reduces the processor overhead during reading and writing cycles.

## Acknowledgements

This work was supported by World Class Institute (WCI) Program of the National Research Foundation of Korea (NRF) Grant Number: WCI 2009-003) and by the Korea Institute of Science and Technology (KIST) grants 2E26190 and 2E26170, and US NIH grants DC005259, NS099691, DC016133 and internal funds from Florida State University. Thanks to J. Verhagen for programming technical assistance.

## Notes

**Conflict of interest** The authors declare no competing financial interests.

