## Supplementary figures and images for "The olfactory bulb contributes to the adaptation of odor responses: the input-output transformation"

### Supplementary Figure 1

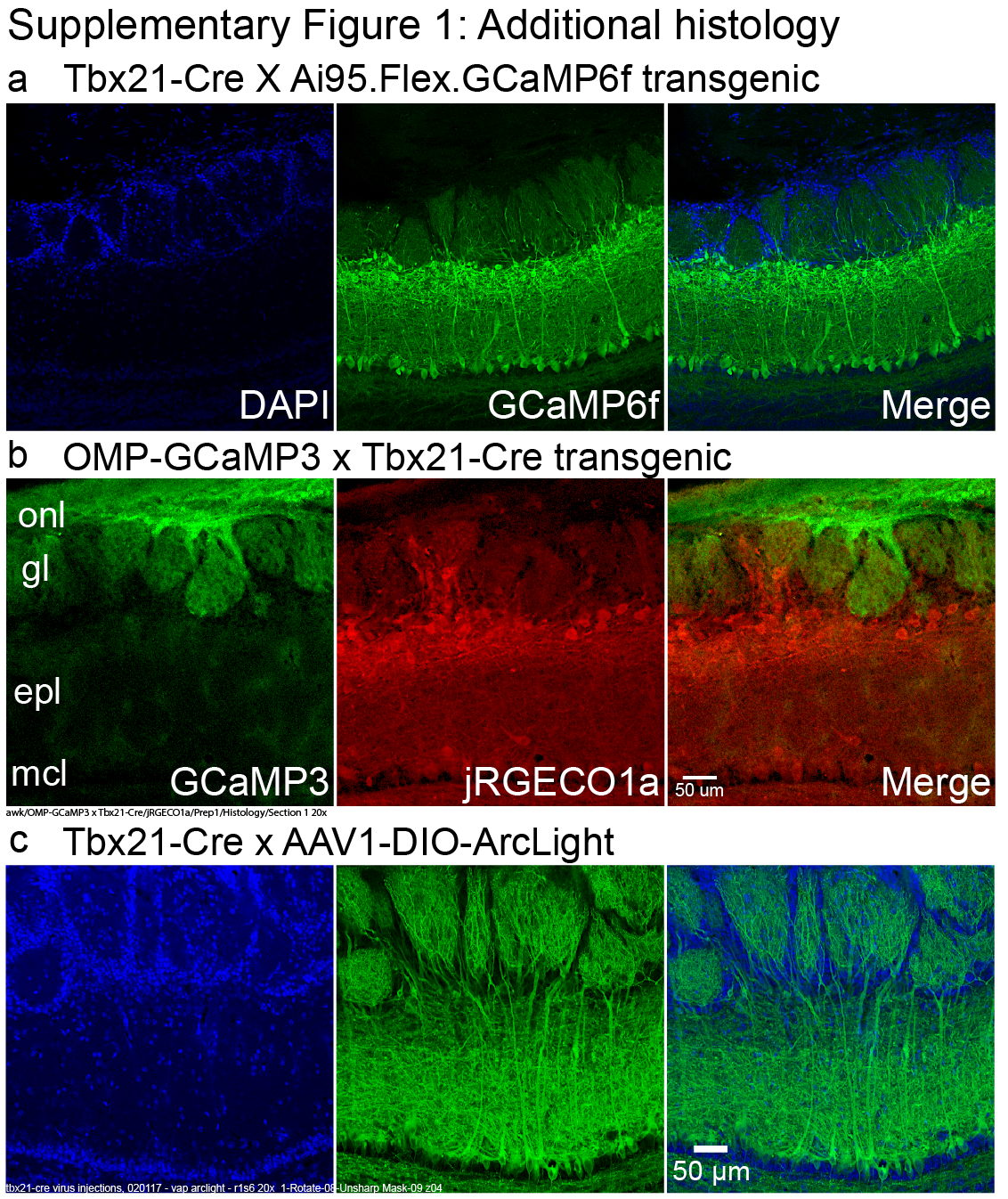

### Supplementary Figure 2

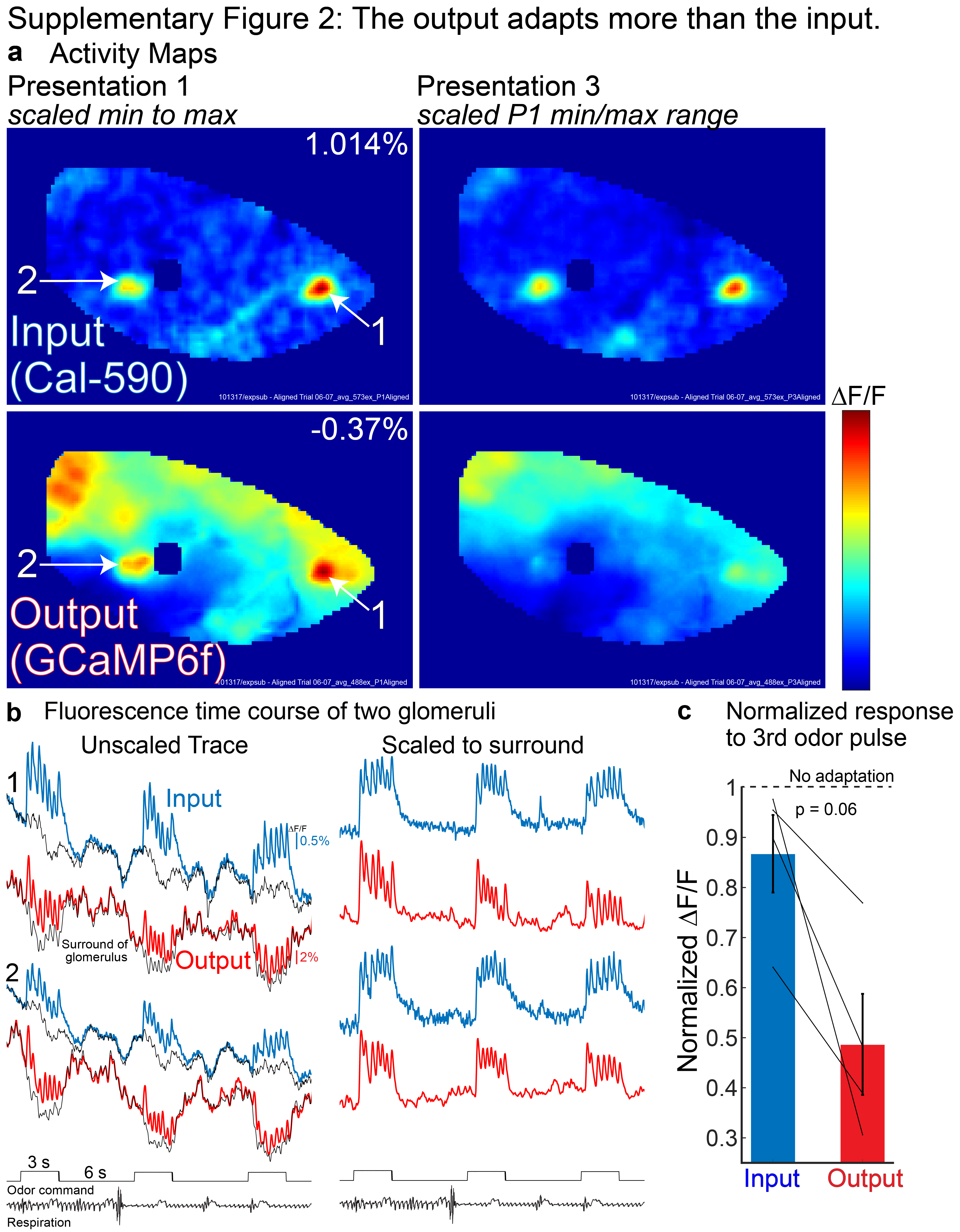

### Supplementary Figure 3

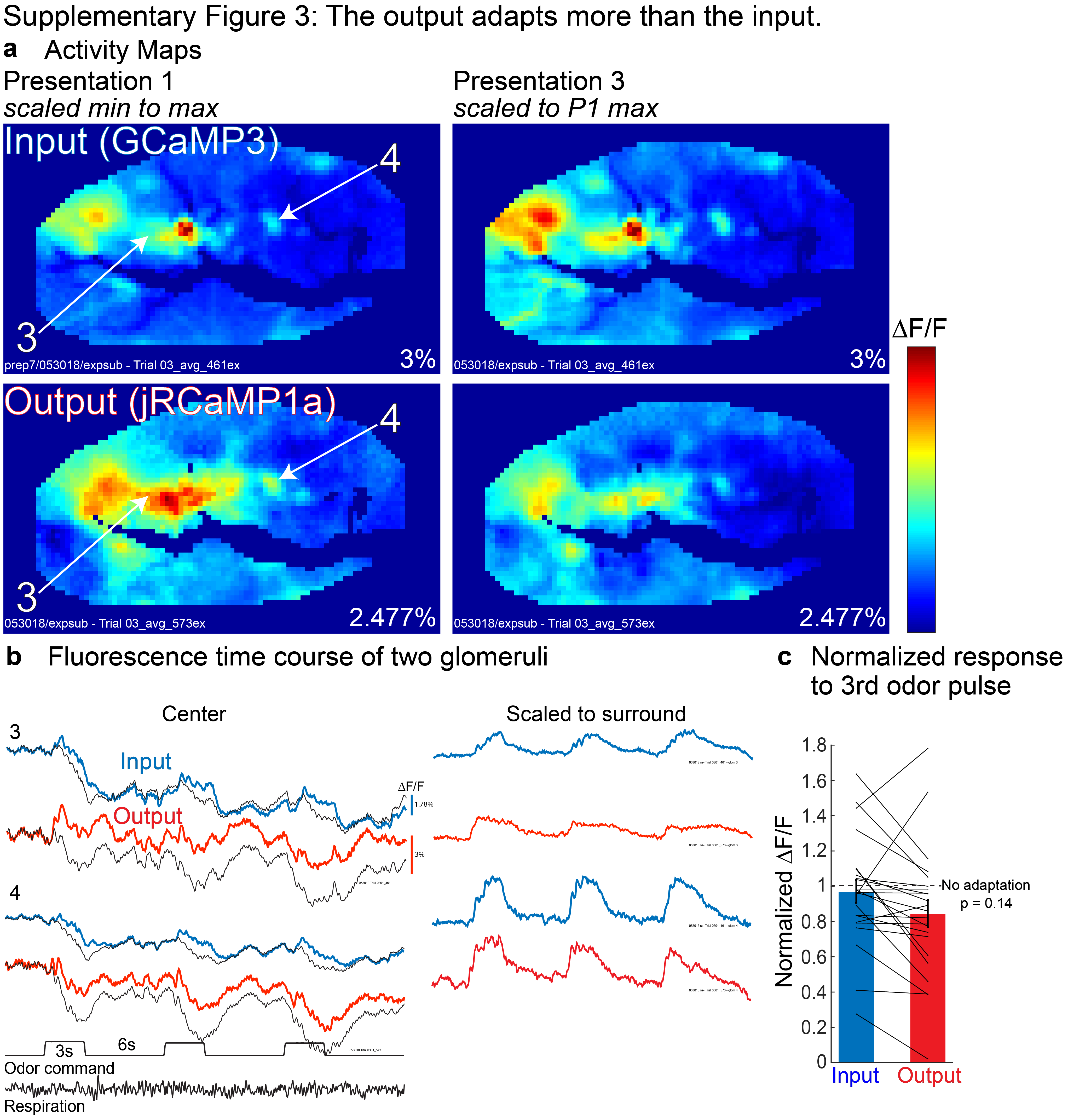

### Supplementary Figure 4

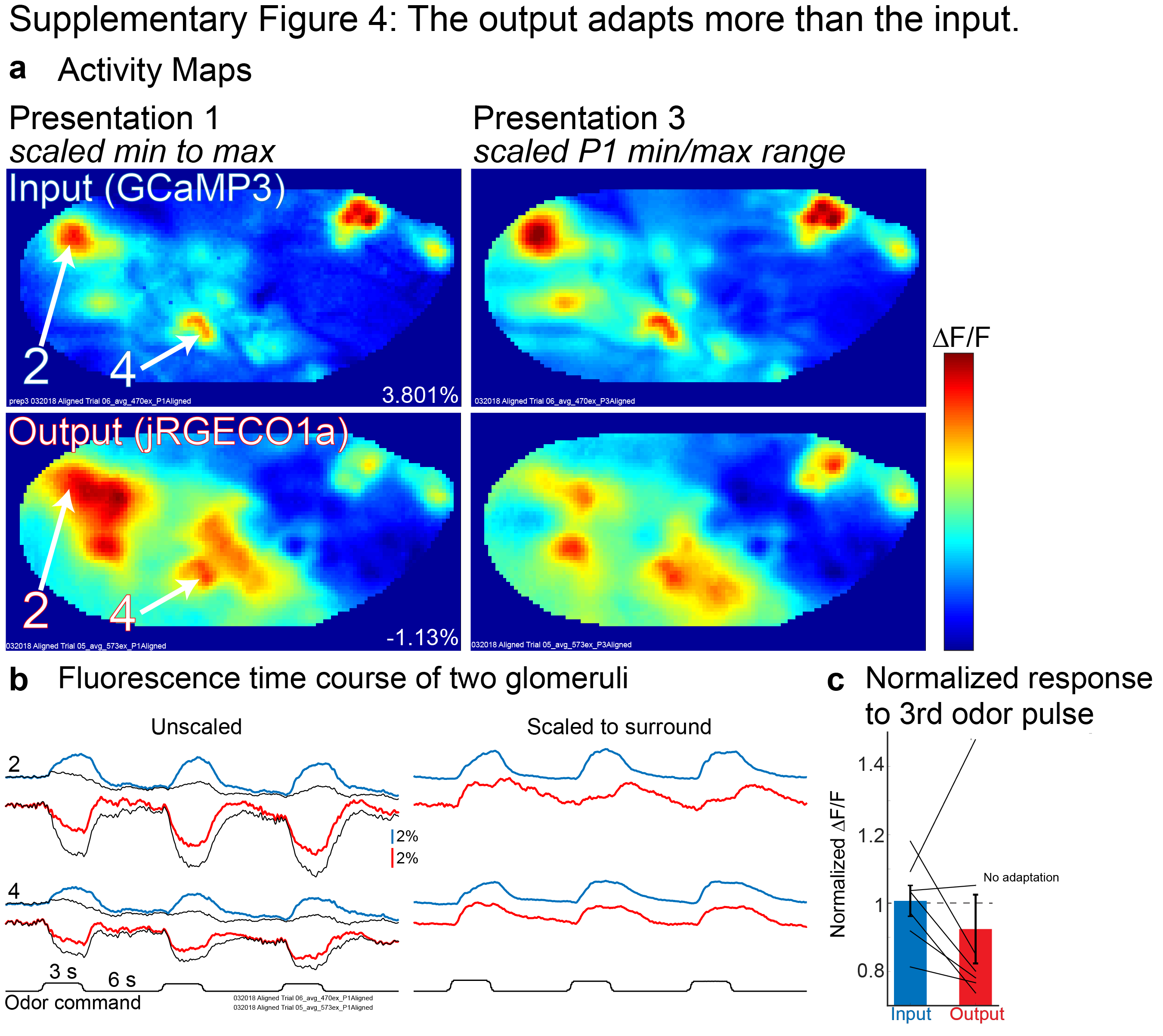

### Supplementary Figure 5

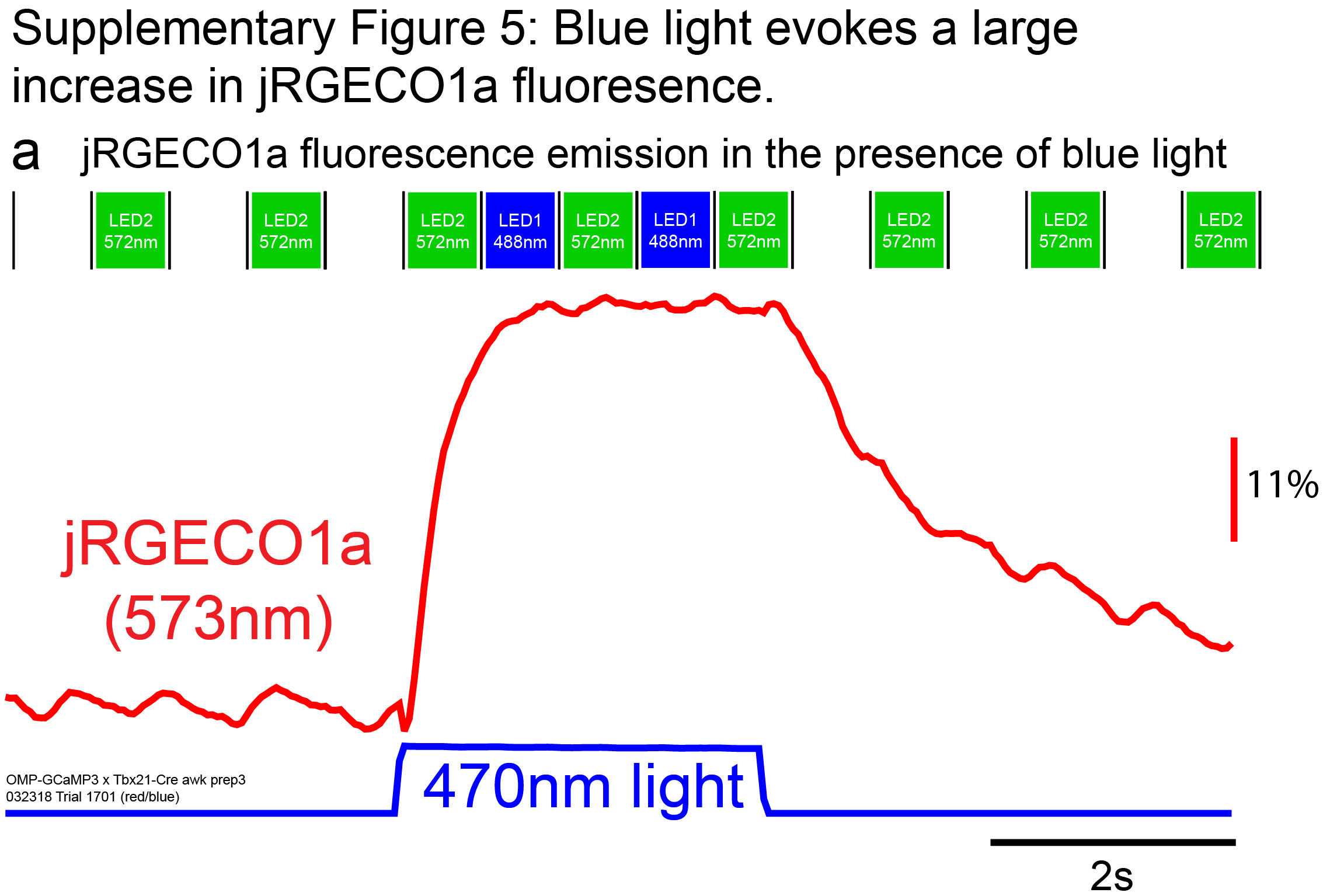

### Supplementary Figure 6

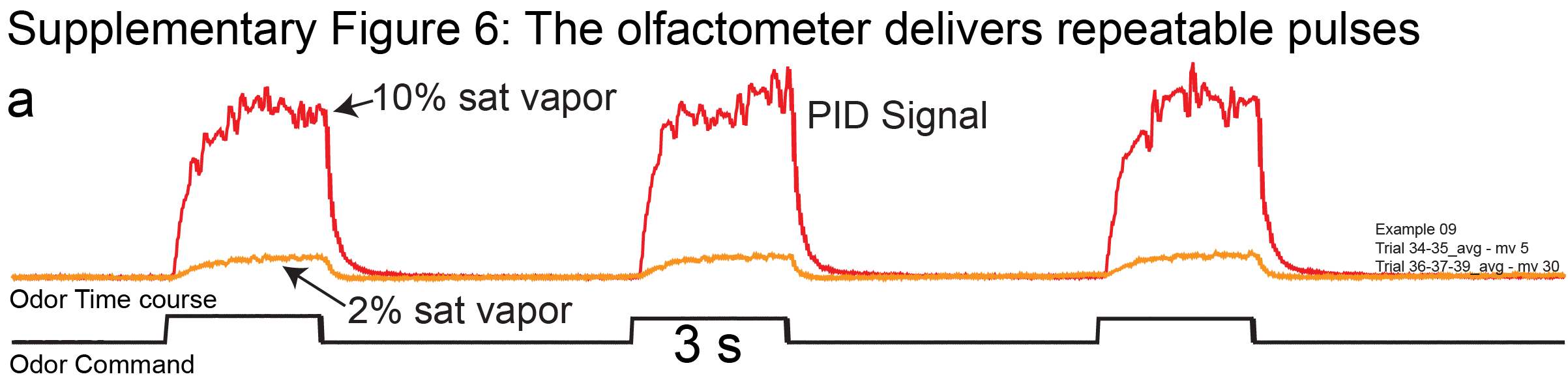
